# Widely naturalized species are not more promiscuous in the use of different nitrogen forms, but benefit more from inorganic nitrogen

**DOI:** 10.1101/2021.08.02.454750

**Authors:** Jianjun Zeng, Yanjie Liu, Mark van Kleunen

## Abstract

Nitrogen (N) is an essential mineral nutrient necessary for plant growth and has been considered a crucial factor influencing invasion success. Former studies mainly focused on responses of alien plants to different levels of N availability. However, in natural soils, N comes in different forms. Few experimental studies have explored responses of naturalized alien species to different N forms, and whether such responses are related to their naturalization success globally.

We selected 22 common herbaceous species native to Germany that have all become naturalized and thus been introduced elsewhere in the world. We grew these species under six different N conditions that differed in the amount or form of N, and assessed their growth performance in a greenhouse experiment.

We found that plants produced more biomass when grown under high N levels than under low N levels, and when N was provided in inorganic form than when provided in organic form. Neither biomass production nor promiscuity to different N forms was related to naturalization success of the species. However, the biomass response to inorganic N, relative to organic N, was stronger for the widely naturalized species than for the less widely naturalized ones.

Our study shows that although the widely naturalized alien species were not more promiscuous than the less widely naturalized species, they took more advantage of the inorganic-N forms. This indicates that naturalization success might be partly driven by a species’ ability to take advantage of increased inorganic N levels.

## 1 Introduction

Many alien plant species that have become naturalized outside of their native ranges pose serious environmental, economic, and human health impacts (Vilà *et al*., 2011; Schaffner *et al*., 2020; Diagne *et al*., 2021). The rise in the number of naturalized alien plant species still shows no sign of saturation (Seebens *et al*., 2017) and is projected to increase from 2005 to 2050 by about 18 % (Seebens *et al*., 2021). Concerns over the impacts of plant invasions have stimulated considerable interest in the mechanisms underlying plant invasions (van Kleunen *et al*., 2010; Funk, 2013; Zhang *et al*., 2020a; Zhang *et al*., 2020b; Liu *et al*., 2021). However, despite considerable progress in our understanding of invasions, it is still unclear why some alien plant species are more successful than others.

The fluctuating-resource hypothesis poses that an alien plant species could establish more easily in a community if the nutrient availability increases (Davis *et al*., 2000). Several studies found support for this hypothesis, and indicate that many successful alien species are opportunists that capitalize more strongly on additional resources than less successful species (Richards *et al*., 2006; Dawson *et al*., 2012b). For example, common alien species showed stronger biomass increases in response to nutrient addition than rare alien species (Dawson *et al*., 2012a). Parepa *et al*. (2013) found that invasive *Fallopia* sp. became more dominant after an increase in nutrients. It was also shown that Central European plant species from more productive habitats are more likely to be invasive elsewhere in the world than species from less productive habitats (Dostál *et al*., 2013). Liu *et al*. 2019 showed that the Central European plants that are widely naturalized elsewhere grow faster than the ones that are not widely naturalized, but they could not related this to differences in nitrogen-acquisition ability of the species. Therefore, more research is needed to explain how some species are able to take more advantage of additional nutrients than others and how this relates to their invasion success.

Nitrogen (N) is an essential mineral nutrient necessary for plant growth, and has been considered a crucial factor influencing invasion success (Seabloom *et al*., 2015; Liu *et al*., 2017a). Nitrogen, however, comes in different forms. Initially, it was thought that only inorganic N, in the form of ammonium (NH_4_^+^) and nitrate (NO_3_^-^) are available to plants, but there is now considerable evidence that many plants also have the ability to take up organic N forms (mainly amino acids, such as glycine and histidine; Näsholm *et al*., 2009; Boudsocq *et al*., 2012; Andersen *et al*., 2017; Liu *et al*., 2017). As a consequence of atmospheric N deposition, but also of fertilizer run-off from agricultural land, not only the overall amount of N has increased, but also the proportions of the soil-available N forms has changed (Krupa, 2003; Song *et al*., 2015). For example, while atmospheric N deposition has resulted in a shift from nitrate-to ammonium-dominance in the United States, the opposite has been observed in China (Liu, X *et al*., 2016; Yu *et al*., 2019). Moreover, due to changes in microbial activity, the proportions of organic to inorganic N might also change, and it was recently shown that in some habitats, such as tundras, amino acids might be the dominant N form (Homyak *et al*., 2021). It is frequently assumed that variation in preferences for various soil N forms could allow plant species to decrease their niche overlap, and to promote coexistence (McKane *et al*., 2002; Ashton *et al*., 2010). Following this logic, a species that can successfully invade, and thus coexist with the resident species, might be more “promiscuous” in the use of different nitrogen forms than less-successful alien species. This would be in line with the long-standing idea that generalists, with a high environmental tolerance, have higher invasion potential than specialists (Baker, 1965; Richards *et al*., 2006).

The preferences for various nitrogen forms are species-specific (Ashton *et al*., 2010; Boudsocq *et al*., 2012; Liu *et al*., 2017; Liu *et al*., 2020). For example, the invasive species *Mikania micrantha, Ipomoea cairica, Wedelia trilobata* and *Bidens pilosa* performed better when supplied with NO_3_^-^ than with NH_4_^+^ (Chen *et al*., 2018). However, the invasive species *Flaveria bidentis* is able to use either NO_3_^-^ or NH_4_^+^ depending on which form is most available in the environments, and this promiscuity probably has contributed to its dominance in many communities (Huangfu *et al*., 2019). To test whether a preference for a certain N form or promiscuity contribute to invasion success in general, we need studies that compare successful with less successful invaders. However, there are many different factors that might determine the invasion success of a species. This variation can be reduced by comparing species that are native to the same region but differ in their invasion success elsewhere (i.e. the source-area approach) (Pyšek *et al*., 2004; van Kleunen *et al*., 2010; Liu *et al*., 2019). Variation can be further reduced by selecting species from similar habitats that are equally common in their native range.

To test whether preferences for certain N forms or promiscuity with regard to N forms could drive variation in naturalization success of alien species, we grew 22 Central European herbaceous grassland species that differ in their global naturalization success under six different N treatments. The six N treatments included one low and five high N availability treatments: 1) low availability with equal amounts of nitrate-nitrogen (NO_3_^-^-N), ammonium nitrogen (NH_4_^+^-N), glycine (Gly-N) and histidine (His-N). 2) high availability with equal amounts of NO_3_^-^-N, NH4^+^-N, Gly-N and His-N, 3) high availability of NO_3_^-^-N only, 4) high availability of NH_4_^+^-N only, 5) high availability of Gly-N only, 6) high availability of His-N only. We compared how biomass and root allocation responses to the N treatments related to the global naturalization success of the species. Specifically, we tested the following questions: (i) Do widely naturalized species overall show a stronger response to increasing N availability than less widely naturalized species? (ii) Do the preferences for the different N forms differ between widely and less widely naturalized species? (iii) Are widely naturalized species overall more promiscuous with regard to the different N forms than less widely naturalized species?

## 2 Materials and Methods

### 2.1 Study species

To increase our ability to generalize the results on N-preference differences among species varying in their global naturalization success (van Kleunen *et al*., 2014), we selected 22 herbaceous plant species from eight families, and with different life spans (Table S1). All 22 species are common natives in Germany (occuring in at least 1928 of the 3000 German grid cells; Table S1), but have become naturalized to different extents in the rest of the world. We determined the number of regions in which each species has become naturalized from the Global Naturalised Alien Flora (GloNAF) database, version 1.2 (van Kleunen *et al*., 2019), which includes data on naturalization success of 13,939 plant species across a total of 1,029 regions. The number of GloNAF regions in which the selected species are recorded as naturalized ranged from 1 to 283 (Table S1). Seeds were either acquired from the field in Konstanz or bought from a commercial seed company (Rieger-Hofmann GmbH, Germany), which produces seeds for grassland-restoration and agricultural purposes (i.e. pastures and meadows; Table S1).

### 2.2 Pre-cultivation and experimental set-up

To test whether the global naturalization success of the study species is related to their N-preferences, we did a greenhouse experiment in the botanical garden of the University of Konstanz (Germany). On 18 and 19 May 2020, seeds were sown into trays (12.0 × 12.0 × 4.5 cm) filled with potting soil (Topferde®, Einheitserde Co., Sinntal-Altengronau, Germany; PH 5.8; 2.0 g/L KCl; 340 mg/L N; 380 mg/L P_2_O_6_; 420 mg/L K_2_O;200 mg/L S; 700 mg/L Mg) for each of the 22 species separatately. We then placed the trays in a greenhouse at a temperature between 22 and 28°C, and a day:night cycle of 16:8 hr.

On 2 June 2020, two weeks after sowing, we selected 18 similarly sized seedlings per species, and transplanted each of them separately into circular 2-L plastic pots (i.e. one individual per pot) filled with a 1:1 nutrient-poor mixture of sand and fine vermiculite. In total, the experiment included 396 pots (i.e., 18 pots per species × 22 species). To test whether responses to the amount of N and the different forms of N vary among the study species and relate to their naturalization success, we assigned the 18 pots of each species to six different N treatments (3 pots per treatment). The N treatments started two weeks after transplanting, and lasted for eight weeks. The six N treatments included one low N availability treatment and five high N availability treatments. Each week, we applied the six different N treatments using six modified Hoagland solutions that only differed from each other in the amount or form of N. In other words, all nutrient solutions contained the same amounts of the other nutrients (e.g. P, K). The low N availability treatment (further referred to as ‘low mixed N’) included a 1:1:1:1 mixture of nitrate-nitrogen (NO_3_^-^-N), ammonium nitrogen (NH_4_^+^-N), glycine (Gly-N) and histidine (His-N). For the five high N availability treatments, the N forms differed: (i) a 1:1:1:1 mixture of NO_3_^-^-N, NH_4_^+^-N, Gly-N and His-N (further referred to as ‘high mixed N’), (ii) only NO^3-^-N, (iii) only NH^4+^-N, (iv) only Gly-N, (v) only His-N. Because the charge of amino acids might determine their diffusion rates in soil (e.g. Homyak *et al*., 2021), we chose glycine, a neutrally charged amino acid, and histidine, a positively charged amino acid. We supplied 40 ml of each of the nutrient solutions to each pot once a week, so that 0.24 mmol and 2.4 mmol of N were provided each time for low mixed N and the five high N availability treatments.

During the whole experiment, we kept all pots at temperatures between 22 and 28°C, and provided supplemental lighting to extend the daily light period to 14 hr. We watered each pot when the soil looked dry, and supplied the same amounts of water to all plants each time.

### 2.3 Measurements

To be able to account for variation in initial sizes of the plants in the analyses, we counted at the start of the experiment the number of true leaves, and measured the length and width of the longest leaf on each plant. Based on these measurements, we calculated a proxy of initial leaf area as the length × width of the largest leaf × the number of true leaves. On 5 August 2020 (i.e. 8 weeks after the start of the N treatments), we assessed survival of the plants, and then started to harvest the ones that had survived. We first harvested the aboveground biomass of each plant, and then washed their roots clean of substrate. The entire belowground biomass harvest took three days, from 6 to 8 August 2020. All aboveground and belowground biomass was dried at 70°C for 72 hours and then weighed. Based on the final aboveground and belowground biomass, we calculated total biomass and root weight ratio (RWR) (belowground biomass/total biomass) for each plant.

### 2.4 Statistics analyses

To test the effects of naturalization success (i.e., the number of regions in which the species naturalized) and the different N treatments on survival, we fitted a binomial generalized linear mixed-effect model using the glmer function in the package “lme4” (Bates *et al*., 2015) in R 4.0.3 (R Core Team, 2013). Similarly, to test the effects of naturalization success and the different N treatments on total biomass and root weight ratio, we fitted linear mixed-effects models using the lme function in the package “nlme” (Pinheiro, 2011). Survival, total biomass and root mass fraction of plants were the response variables in the models. To meet the normality assumption, total biomass was cubic-transformed. We included naturalization success (i.e. the number of naturalized regions per species), the different N treatments (i.e. low mixed N, high mixed N, high NO_3_^-^-N, high NH_4_^+^-N, high Gly-N and high His-N) and their interactions as fixed effects in the models. We did not include the interactions in the model analysing survival, because inclusion resulted in model-convergence failure. To test more specifically which of the N treatments differed from each other, we created five orthogonal contrasts by coding dummy variables (Schielzeth, 2010): low mixed N *vs* the average of all high N treatments (N_low-high_), high mixed N *vs* the average of the non-mixed high N treatments (N_mix-nonmix_), the average of the two high inorganic N treatments *vs* the average of the two high organic N treatments (N_inorganic-organic_), high NO_3_^-^-N *vs* high NH_4_^+^-N (N_NO3-NH4_), high Gly-N *vs* high His-N (N_gly-his_). Because variation in initial plant size might contribute to differences in final plant performance, we also added initial leaf area as a scaled naturalLlogLtransformed covariate in the models. To account for phylogenetic non-independence of species and for non-independence of replicates of the same species, we included species nested within family as random factors in all models. As the homoscedasticity assumption was violated in linear mixed models, we included variance structures in those models to allow different variances per species using the function “varIdent” in the R package nlme (Pinheiro, 2011). In all models, we assessed the significance of naturalization success, N treatments (and its contrasts), and their interactions with likelihood-ratio tests (Zuur *et al*., 2009).

To assess whether promiscuity to the different N forms was related to naturalization success of the species, we needed an index indicating whether the biomass responses of a species to the different N forms was relatively constant (indicating promiscuity) or very variable. We therefore first calculated for each species in each of the five high N treatments the log-response ratio of the average biomass in the respective high N treatment vs the average biomass of the species in the low N treatment. Then as measure of how variable this response was, we calculated the standard deviation across the five log-response ratios of each species. Finally, to test whether the standard deviation of the log-response ratios was related to naturalization success of the species, we calculated Pearson’s correlation coefficient.

## 3 Results

Of the initial 396 plants, 50 (12.6%) had died by the end of the experiment. Survival was not related to naturalization success of the species, but was significantly affected by the N treatments (Table S2). Overall, survival was lower in the high N treatments than in the low N treatment, and was lowest when plants received N in organic form as glycine (Table S2, Fig. S1).

Among the surviving plants, and averaged across the six N treatments, total biomass was not related to naturalization success of the species (Table 1; Fig. 1A). Averaged across all species, plants produced more biomass (+159.7%) when grown under high N levels than under low N levels (Fig. 1A, B). Among the high non-mixed N treatments, plants also produced more total biomass (+73.4%) when N was provided in inorganic form than when provided in organic form (Table 1; Fig. 1A, B). This increase in biomass production in response to inorganic vs organic N was larger for the widely naturalized species than for the less widely naturalized species (significant Nat × N_inorganic-organic_ in Table 1, Fig. 1A).

**Table 1.** Results of linear mixed effects models testing the effects of naturalization success (number of regions where the species is naturalized), nitrogen treatment and their interaction on total biomass (cubic-root-transformed) and root weight ratio. Initial leaf area was included as a covariable, and for the N treatment, we made five planned orthogonal contrasts. For the random terms, the standard deviations (SD) are provided.

| Terms | df | Cbrt (total biomass) |  | Root weight ratio |  |
| --- | --- | --- | --- | --- | --- |
|  |  | X<sup>2</sup> | P | X<sup>2</sup> | P |
| *Fixed terms* |  |  |  |  |  |
| Initial leaf area | 1 | 3.00 | 0.083 | 8.41 | **0.004** |
| Naturalization success (Nat) | 1 | 0.17 | 0.679 | 0.21 | 0.644 |
| N treatment (N) | 5 | 130.26 | **<0.001** | 99.35 | **<0.001** |
| N<sub>low-high</sub> | 1 | 50.30 | **<0.001** | 97.45 | **<0.001** |
| N<sub>mix-nonmix</sub> | 1 | 1.43 | 0.232 | 0.10 | 0.747 |
| N<sub>inorganic-organic</sub> | 1 | 74.54 | **<0.001** | 0.16 | 0.692 |
| N<sub>NO3-NH4</sub> | 1 | 2.21 | 0.137 | 0.92 | 0.337 |
| N<sub>gly-his</sub> | 1 | 1.89 | 0.170 | 0.67 | 0.414 |
| Nat × N | 5 | 9.69 | 0.084 | 13.85 | **0.017** |
| Nat × N<sub>low-high</sub> | 1 | 0.41 | 0.522 | 2.63 | 0.105 |
| Nat × N<sub>mix-nonmix</sub> | 1 | 0.44 | 0.506 | 3.89 | **0.049** |
| Nat × N<sub>inorganic-organic</sub> | 1 | 4.31 | **0.040** | 1.52 | 0.217 |
| Nat × N<sub>NO3-NH4</sub> | 1 | 0.70 | 0.404 | 1.64 | 0.201 |
| Nat × N<sub>gly-his</sub> | 1 | 3.85 | 0.050 | 4.17 | **0.041** |
| *Random terms* |  |  |  |  |  |
|  |  | SD |  | SD |  |
| Family |  | 0.1782 |  | 0.05302 |  |
| Species* |  | 0.3004 |  | 0.05029 |  |
| Residual |  | 0.2878 |  | 0.13709 |  |

\*Each species was allowed to have its own variance, and the standard deviations shown are those for *Daucus carota*. The multiplication factors to calculate the standard deviations of the other species are provided in Table S3.

**Figure 1.**
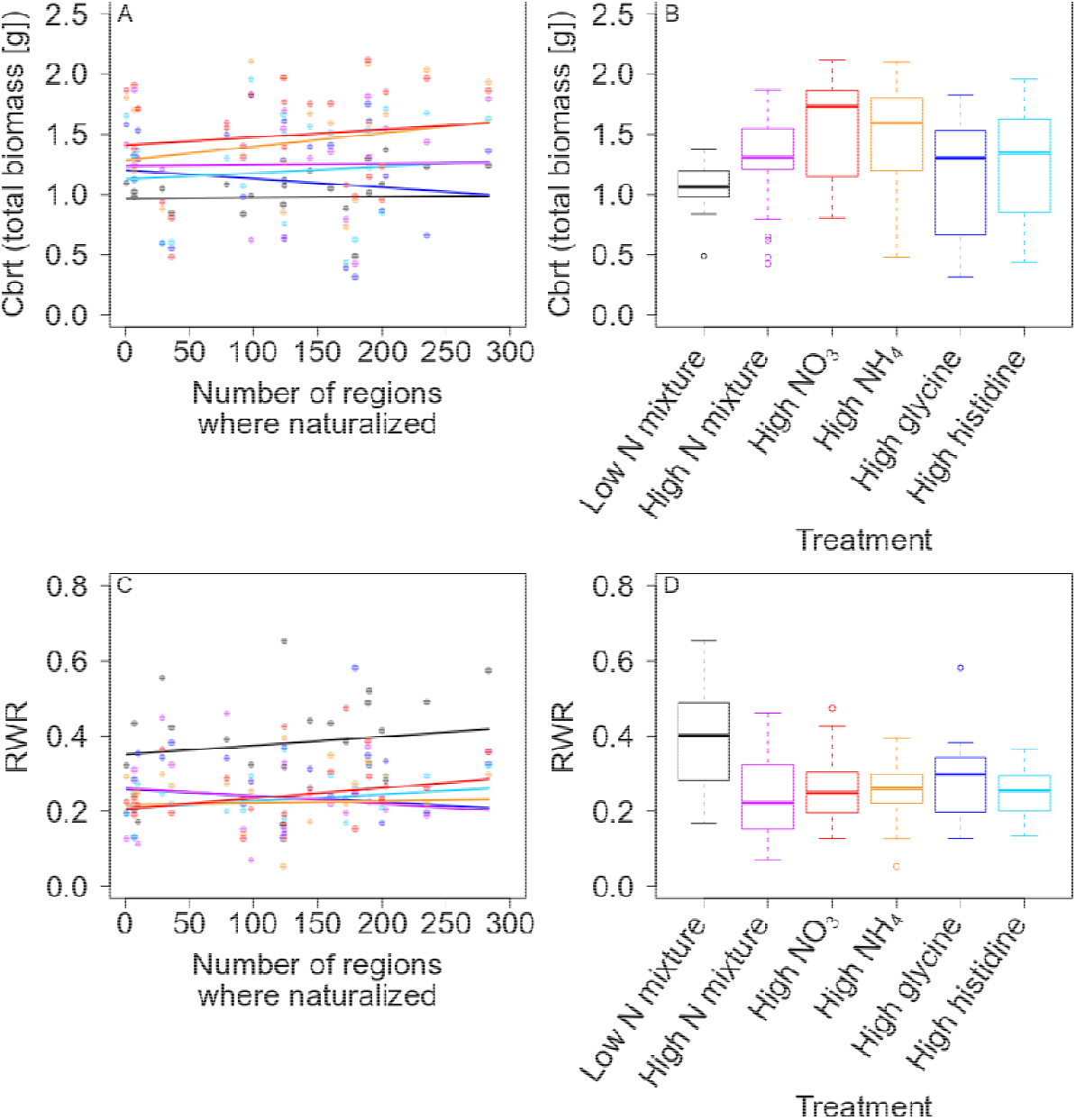
Total biomass (cubic transformed; A, B) and root weight ratio (RWR; C,D) of the 22 study species in response to the one low and five high N treatments. A and C are scatterplots of the species mean values of the measured traits vs the naturalization success of the species, with fitted relationships from the linear mixed models. B and D are boxplots showing the distribution of the species mean values for each of the N treatments. The boxes indicate the interquartile range around the median (thick horizontal line), whiskers extend to 1.5 times the interquartile range, and open circles indicate outliers. The colors in A and C correspond to the treatment colors in B and D.

The root weight ratio (RWR) of plants was also not related to naturalization success of the species (Fig. 1C), and was decreased (−33.1%) when plants were grown at high N levels (Fig. 1C, D). Averaged across all species, there were no significant differences in RWR among the different high N treatments (Table 1; Fig. 1C, D). However, there was a significant interaction between the naturalization extent and the N treatment (Table 1). While RWR decreased with increasing naturalization success for the high N mixture treatment and the high glycine treatment, it increased for the other N treatments (Table 1, Fig. 1C).

The species’ standard deviation for the log-response ratios of biomass in the high N treatments vs biomass in the low N treatment was not significantly related to the naturalization success of the species (r = 0.219, n = 22, *P* = 0.328; Fig. 2). In other words, promiscuity to the different N forms was not related to naturalization success.

**Figure 2.**
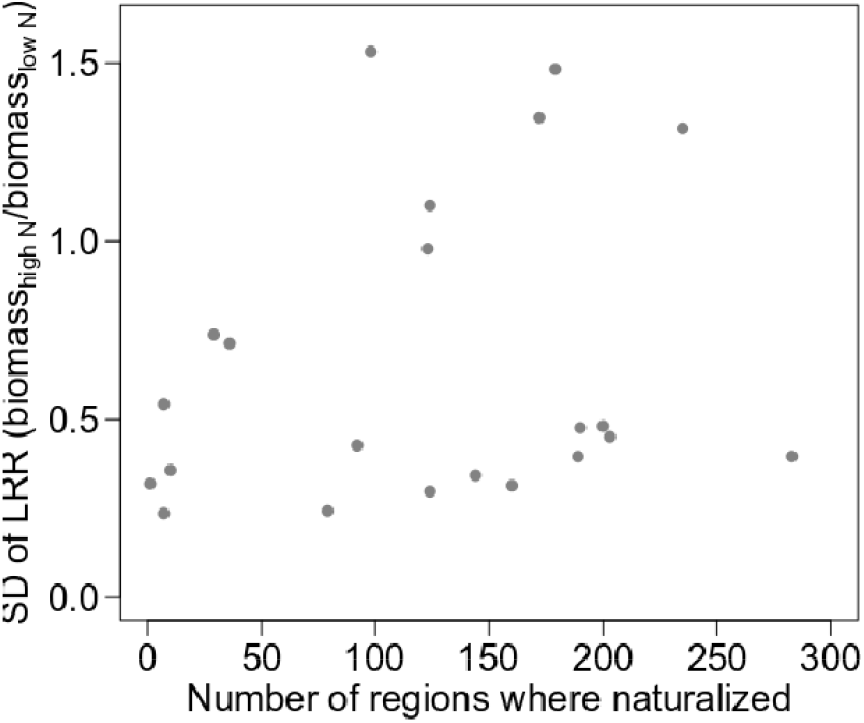
Scatterplot of the standard deviation (SD) of the log response ratios (LRR) of the five different high N treatments relative to the low N treatment of each species vs the naturalization success of the species. A low SD would indicate that a species would respond similarly to the different N forms (i.e. that it would be promiscuous). The correlation was not significant (r = 0.219, n = 22, P = 0.328).

## 4 Discussion

We tested whether preferences for and promiscuity to different N forms of Central European plant species was related to their naturalization success elsewhere. Not surprisingly, plants increased their performance when they had access to more N. However, we found that they benefited more from additional inorganic N than from organic N, and that high levels of glycine even increased mortality. Neither average biomass production nor promiscuity to different N forms was related to naturalization success of the species. However, the biomass response to inorganic N, relative to organic N, was stronger for the widely naturalized species than for the less widely naturalized ones. This indicates that naturalization success might be partly driven by a species’ ability to take advantage of increased inorganic N.

Overall, plants increased their biomass production when growing under high N-conditions relative to when grown under low N. This is not surprising because N availability is generally a limiting factor for plant growth (LeBauer & Treseder, 2008). We also found that plants allocated less biomass to their root systems under high N conditions than under low N. This is in line with the predictions of resource-limitation theory, which states that plant species should invest relatively more biomass into those plant structures that allow them to acquire more of the most limiting resource (Bloom *et al*., 1985; Poorter *et al*., 2012; Liu *et al*., 2016; Liu *et al*., 2019).

Numerous studies have shown that species that are more successful, either as natives or aliens, take more advantage of additional nutrients (Dawson *et al*., 2012b; Parepa *et al*., 2013; Liu *et al*., 2017b). However, here, we did not find evidence that the biomass or root-allocation responses to high N availability were associated with the global naturalization success of the study species. This corroborates one of our previous studies, which showed that global naturalization success of Central European plants is not related to their N-acquisition ability (Liu *et al*., 2019). However, although we did not find a significant relationship between naturalization success and the overall response to an increased N availability, we found that biomass of the widely naturalized species increased more strongly in response to the inorganic N forms than was the case for the less widely naturalized species.

Evidence is accumulating that many species are able to take up nitrogen in organic form (Ashton *et al*., 2010; Boudsocq *et al*., 2012; Liu *et al*., 2017). Indeed, we found that overall species increased their biomass when they were provided with the amino acids glycine or histidine instead of a low-N mixture. Nevertheless, plants benefited more from inorganic nitrogen (NO_3_^-^, NH_4_^+^) than from organic N. The latter is in line with previous studies, and might indicate that inorganic-N is the dominant N form in nature to which most species have adapted (Liu *et al*., 2017). We cannot fully exclude the possibility that some of the organic N in our treatments was mineralized, and that the plants took up the resulting inorganic N. However, if that would have been the case, we would expect that all species would have benefited from organic N addition and subsequent mineralization. Because there were also several species that produced less biomass in response to organic N additon (Fig. 1A, B), and because survival was lowest in the high glycine treatment, it seems likely that the species that benefited took up the organic N forms directly.

As mentioned above the widely naturalized species benefited more from inorganic N, relative to organic N, than less widely naturalized species. In particular, the widely naturalized plants tended to benefit less from the addition of glycine, relative to histidine (marginally significant Nat × N_gly-his_ interaction in Table 1, *P* = 0.050; Fig. 1A). Overall, these results suggest that there is a trade off between species’ responses to inorganic and organic N forms, and that those that benefit the most from inorganic N have the highest naturalization potential.

Previous studies have shown that the different inorganic-N forms, NO_3_^-^ and NH_4_^+^, could affect plant performance differently (Ashton *et al*., 2010; Huangfu *et al*., 2016; Chen *et al*., 2018; Liu *et al*., 2020). We found that performance of our study species overall did not significantly differ between the NO_3_^-^ and NH_4_^+^ treatments. Therefore, our findings suggest that changes in the NO_3_^-^/NH_4_^+^ ratio induced by atmospheric N deposition (Boxman *et al*., 2008) might not affect the overall performance of the plants.

In all the five high N treatments, plants invested on average less biomass in their roots than in the low N treatment. Averaged across species, root weight ratios did not differ among the different N treatments. However, we found that in the high N mixture and high glycine treatments, the relative biomass investment into roots declined with the extent of naturalization of the species, while the reverse was true for the other N treatments. The reasons for this are unknown. However, the fact that the widely naturalized species produced low root weight ratios in the high glycine treatment might have contributed to their low biomass production in this treatment. On the other hand, the widely naturalized species invested more biomass in roots, and also produced more biomass than the less widely naturalized species in the high histidine treatment. This shows that plants do not benefit equally from all organic N forms, and this might be related to the charges of the amino acids (Homyak *et al*., 2021). Although glycine is often considered to be a common amino acid in soils, the same can be true for histidine, as well as for amino acids that we did not included in our experiment (e.g. lysine, arginine and serine) (Lipson & Näsholm, 2001; Tian *et al*., 2020). Future studies on N responses of plant species differing in their naturalization success should also consider these other amino acids as well as other organic N forms that could be taken up by some plants, such as quaternary ammonium compounds (Warren, 2013).

## 5 Conclusions

Our study is the first multi-species experiment, using the source-area approach, to compare the preferences for specific N forms and promiscuity to different N forms among species that differ in their global naturalization success. While the widely naturalized alien species were not more promiscuous than the less widely naturalized species, they took more advantage of the inorganic-N forms. As these forms have globally increased due to atmospheric nitrogen deposition, our results suggest that one does not have to be a generalist in order to become widely naturalized around the world. Instead, it helps to specialize in the most common forms of N available to plants.

## Acknowledgments

We thank Otmar Ficht, Maximilian Fuchs, Duo Chen, Guanwen Wei and Beate Rüter forpracical assistance. J.Z thanks the funding from China Scholarship Council. Y.L acknowledges funding from Chinese Academy of Sciences (Y9B7041001).

## Author contributions

M.v.K conceived the idea, M.v.K, J.Z. and Y.L designed the experiment. J.Z. performed the experiment and collected the data. J. Z., M.v.K and Y.L. analyzed the data and wrote the manuscript.

## Data and code accessibility

Should the manuscript be accepted, the data supporting the results will be archived in Dryad and the data DOI will be included at the end of the article.

## Supplementary materials

### Method S1 Recipes for the nutrient solutions used in the experiment

We prepared the modified Hoagland nutrient-stock solutions according to the recipes given below. Before each round of fertilizer application, we used the stock-solution amounts indicated per 1 liter of nutrient solution:

#### High NO_3_^□^-N treatment

1. 1.00 M·L^-1^ Ca(NO_3_)_2_: 30 ml
2. 1.00 M·L^-1^ KH_2_PO_4_: 4 ml
3. 1.00 M·L^-1^ KCl: 20 ml
4. 1.00 M·L^-1^ Ca(Cl)_2_: 20 ml
5. 1.00 M·L^-1^ MgSO_4_: 8 ml
6. 1000 mg/liter iron from iron chelate (Fe-EDTA): 10 ml

#### High NH_4_^+^-N treatment

1. 1.00 M·L^-1^ (NH_4_)_2_SO_4_: 30 ml
2. 1.00 M·L^-1^ KH_2_PO_4_: 4 ml
3. 1.00 M·L^-1^ KCl: 20 ml
4. 1.00 M·L^-1^ Ca(Cl)_2_: 20 ml
5. 1.00 M·L^-1^ MgSO_4_: 8 ml
6. 1000 mg/liter iron from iron chelate (Fe-EDTA): 10 ml

#### High glycine treatment

1. 1.00 M·L^-1^ glycine (mol wt = 75.07, 1 N atom): 60 ml
2. 1.00 M·L^-1^ KH_2_PO_4_: 4 ml
3. 1.00 M·L^-1^ KCl: 20 ml
4. 1.00 M·L^-1^ Ca(Cl)_2_: 20 ml
5. 1.00 M·L^-1^ MgSO_4_: 8 ml
6. 1000 mg/liter iron from iron chelate (Fe-EDTA): 10 ml

#### High histidine treatment

1. 0.25 M·L^-1^ L-histidine (mol wt = 155.15, 3 N atoms): 80 ml
2. 1.00 M·L^-1^ KH_2_PO_4_: 4 ml
3. 1.00 M·L^-1^ KCl: 20 ml
4. 1.00 M·L^-1^ Ca(Cl)_2_: 20 ml
5. 1.00 M·L^-1^ MgSO_4_: 8 ml
6. 1000 mg/liter iron from iron chelate (Fe-EDTA): 10 ml

#### High N-mixture treatment

1. 1.00 M·L^-1^Ca(NO_3_)_2_: 7.5 ml
2. 1.00 M·L^-1^ (NH_4_)_2_SO_4_: 7.5 ml
3. 1.00 M·L^-1^glycine (mol wt = 75.07, 1 N atom): 15 ml
4. 0.25 M·L^-1^ L-histidine (mol wt = 155.15, 3 N atoms): 20 ml
5. 1.00 M·L^-1^ KH_2_PO_4_: 4 ml
6. 1.00 M·L^-1^ KCl: 20 ml
7. 1.00M·L^-1^ Ca(Cl)_2_: 20 ml
8. 1.00M·L^-1^ MgSO_4_: 8 ml
9. 1000 mg/liter iron from iron chelate (Fe-EDTA): 10 ml

#### Low N-mixture treatment

1. 0.10 M·L^-1^Ca(NO_3_)_2_: 7.5 ml
2. 0.10 M·L^-1^ (NH_4_)_2_SO_4_: 7.5 ml
3. 0.10 M·L^-1^glycine (mol wt = 75.07, 1 N atom): 15 ml
4. 0.10 M·L^-1^ L-histidine (mol wt = 155.15, 3 N atoms): 5 ml
5. 1.00 M·L^-1^ KH_2_PO_4_: 4 ml
6. 1.00 M·L^-1^ KCl: 20 ml
7. 1.00 M·L^-1^ Ca(Cl)_2_: 20 ml
8. 1.00 M·L^-1^ MgSO_4_: 8 ml
9. 1000 mg/liter iron from iron chelate (Fe-EDTA): 10 ml

In addition, we added per liter of nutrient solution 4 ml of a micronutrient-stock solution, which contained per liter:

2.86 g H_3_BO_3_

1.81 g MnCl_2_·4H_2_O

0.22 g ZnSO_4_·7H_2_O

0.08 g CuSO_4_·5H_2_O

0.02 g H_2_MoO_4_·H_2_O (assaying 85% MoO_3_)

**Table S1.** Information on the the number of regions where the species is known to be naturalized, sowing time and life span of the 22 study cies. The species are grouped per family.

| Family | Species | Sowing date<br>(dd.mm.yy) | Life span<sup>1</sup> | Occupied grid cells<br>in Germany<sup>2</sup> | Naturalized<br>regions<sup>3</sup> |
| --- | --- | --- | --- | --- | --- |
| Apiaceae | *Daucus carota* L. | 18.05.20 | Biennial-Perennial | 2942 | 172 |
| Asteraceae | *Picris hieracioides* L. | 18.05.20 | Biennial-Perennial | 2074 | 29 |
| Asteraceae | *Artemisia vulgaris* L. | 19.05.20 | Perennial | 2965 | 98 |
| Asteraceae | *Hypochaeris radicata* L. | 18.05.20 | Perennial | 2917 | 190 |
| Brassicaceae | *Lepidium campestre* (L.) R. Br.* | 19.05.20 | Biennial | 1928 | 92 |
| Caprifoliaceae | *Knautia arvensis* (L.) Coult. | 19.05.20 | Perennial | 2855 | 36 |
| Lamiaceae | *Scutellaria galericulata* L. | 18.05.20 | Perennial | 2749 | 1 |
| Leguminosae | *Ononis spinosa* L. | 18.05.20 | Perennial | 2553 | 10 |
| Leguminosae | *Lotus corniculatus* L. | 18.05.20 | Annual-Biennial-Perennial | 2931 | 124 |
| Leguminosae | *Trifolium pratense* L.* | 18.05.20 | Annual-Biennial-Perennial | 2989 | 179 |
| Leguminosae | *Medicago lupulina* L. | 18.05.20 | Annual-Biennial-Perennial | 2969 | 203 |
| Poaceae | *Poa angustifolia* L. | 18.05.20 | Perennial | 2050 | 7 |
| Poaceae | *Bromus sterilis* L. | 19.05.20 | Annual-Biennial | 2651 | 79 |
| Poaceae | *Agrostis gigantea* Roth. | 18.05.20 | Perennial | 2108 | 124 |
| Poaceae | *Phleum pratense* L. | 18.05.20 | Perennial | 2978 | 144 |
| Poaceae | *Bromus hordeaceus* L. | 18.05.20 | Annual-Biennial-Perennial | 2928 | 160 |
| Poaceae | *Holcus lanatus* L. | 18.05.20 | Perennial | 2975 | 189 |
| Poaceae | *Poa pratensis* L. | 18.05.20 | Perennial | 2876 | 200 |
| Poaceae | *Dactylis glomerata* L. | 18.05.20 | Perennial | 2910 | 235 |
| Poaceae | *Lolium perenne* L. | 18.05.20 | Perennial | 2981 | 283 |
| Scrophulariaceae | *Verbascum densiflorum* Bertol. | 18.05.20 | Biennial | 2073 | 7 |
| Scrophulariaceae | *Verbascum thapsus* L. | 18.05.20 | Biennial | 2564 | 123 |

<sup>1</sup>The data were obtained from the BioFlor database (http://www2.ufz.de/bioflor/index.jsp) and Wikipedia (https://www.wikipedia.org/).
<sup>2</sup>The data on grid cell frequency (n = 3000) in Germany were obtained from the FloraWeb database (https://www.floraweb.de/).
<sup>3</sup>The number of naturalized regions per species were obtained from the GloNAF database (van Kleunen *et al.*, 2019).
\*Seeds of the species marked with asterisk were collected from the field in Konstanz. Seeds of the other species were acquired from Rieger-Hofmann GmbH, Germany.

**Table S2.**
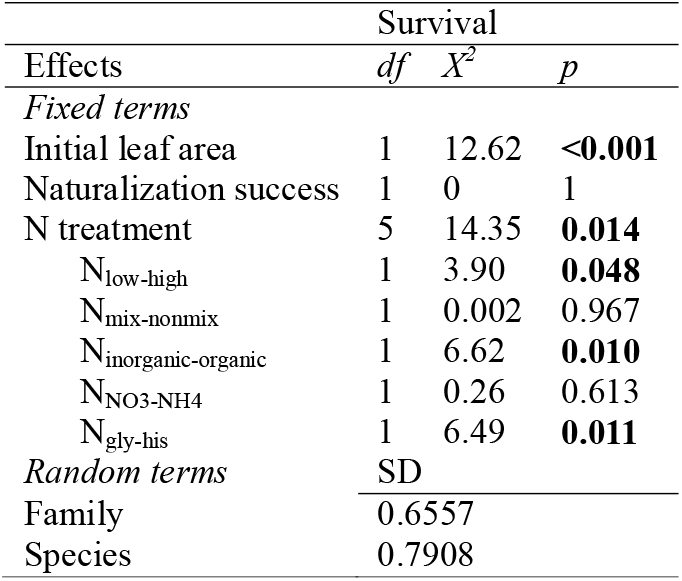
Results of binomial generalized linear mixed effects models testing the effects of naturalization success (number of regions where the species is naturalized) and nitrogen treatment on survival of the plants. Initial leaf area was included as a covariable, and for the N treatment, we made five planned orthogonal contrasts. For the random terms, the standard deviations (SD) are provided.

| Effects | Survival |  |  |
| --- | --- | --- | --- |
| | *df* | $X^2$ | *p* |
| *Fixed terms* |  |  |  |
| Initial leaf area | 1 | 12.62 | **<0.001** |
| Naturalization success | 1 | 0 | 1 |
| N treatment | 5 | 14.35 | **0.014** |
| N<sub>low-high</sub> | 1 | 3.90 | **0.048** |
| N<sub>mix-nonmix</sub> | 1 | 0.002 | 0.967 |
| N<sub>inorganic-organic</sub> | 1 | 6.62 | **0.010** |
| N<sub>NO3-NH4</sub> | 1 | 0.26 | 0.613 |
| N<sub>gly-his</sub> | 1 | 6.49 | **0.011** |
| *Random terms* | SD |  |  |
| Family | 0.6557 |  |  |
| Species | 0.7908 |  |  |

**Table S3.** Multiplication factors to derive the standard deviation for each of the species, except for *Daucus carota*, for the linear mixed model results shown in Table 1. To calculate the standard deviation of each species, its multiplication factor should be multiplied with the species standar deviation shown in Table 1.

| Species | Total biomass | RWR |
| --- | --- | --- |
| *Artemisia vulgaris* | 1.51162 | 0.54320 |
| *Hypochaeris radicata* | 1.15657 | 0.78429 |
| *Picris hieracioides* | 1.08179 | 0.71406 |
| *Lepidium campestre* | 0.59368 | 0.51003 |
| *Knautia arvensis* | 0.92849 | 0.50488 |
| *Scutellaria galericulata* | 0.91962 | 0.72244 |
| *Lotus corniculatus* | 0.96557 | 0.35055 |
| *Medicago lupulina* | 0.94659 | 0.63884 |
| *Ononis spinosa* | 0.68986 | 0.83549 |
| *Trifolium pratense* | 0.56547 | 1.05839 |
| *Agrostis gigantea* | 0.82138 | 0.60249 |
| *Bromus hordeaceus* | 0.45783 | 0.41217 |
| *Bromus sterilis* | 0.64543 | 0.79103 |
| *Dactylis glomerata* | 1.09540 | 0.64362 |
| *Holcus lanatus* | 0.66090 | 0.65678 |
| *Lolium perenne* | 0.89267 | 0.68653 |
| *Phleum pratense* | 0.70983 | 0.63901 |
| *Poa angustifolia* | 0.68921 | 0.27115 |
| *Poa pratensis* | 1.00356 | 0.44185 |
| *Verbascum densiflorum* | 1.12400 | 0.62871 |
| *Verbascum thapsus* | 1.50388 | 0.53096 |

**Figure S1.**
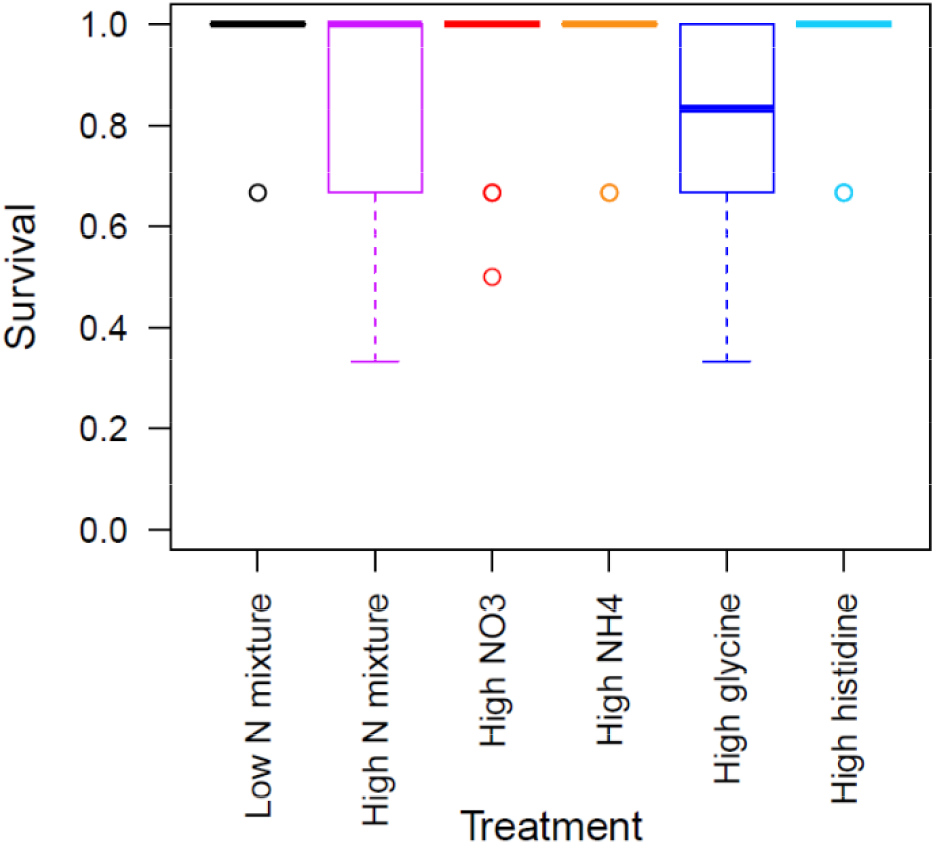
Boxplots of survival of the plants in response to the one low and five high N treatments. The boxes indicate the interquartile range around the median (thick horizontal line), whiskers extend to 1.5 times the interquartile range, and open circles indicate outliers.

